# Genomics-Based Identification of Microorganisms in Human Ocular Body Fluid

**DOI:** 10.1101/176529

**Authors:** Philipp Kirstahler, Søren Solborg Bjerrum, Alice Friis-Møller, Morten la Cour, Frank M. Aarestrup, Henrik Westh, Sünje Johanna Pamp

## Abstract

Advances in genomics have the potential to revolutionize clinical diagnostics. Here, we examine the microbiome of vitreous (intraocular body fluid) from patients who developed endophthalmitis following cataract surgery or intravitreal injection. Endophthalmitis is an inflammation of the intraocular cavity and can lead to a permanent loss of vision. As controls, we included vitreous from endophthalmitis-negative patients, balanced salt solution used during vitrectomy, and DNA extraction blanks. We compared two DNA isolation procedures and found that an ultraclean production of reagents appeared to reduce background DNA in these low microbial biomass samples. We created a curated microbial genome database (>5700 genomes) and designed a metagenomics workflow with filtering steps to reduce DNA sequences originating from: i) human hosts, ii) ambiguousness/contaminants in public microbial reference genomes, and iii) the environment. Our metagenomic read classification revealed in nearly all cases the same microorganism than was determined in cultivation‐ and mass spectrometry-based analyses. For some patients, we identified the sequence type of the microorganism and antibiotic resistance genes through analyses of whole genome sequence (WGS) assemblies of isolates and metagenomic assemblies. Together, we conclude that genomics-based analyses of human ocular body fluid specimens can provide actionable information relevant to infectious disease management.

## Introduction

Genomics-based analyses of patient specimens have the potential to provide actionable information that could facilitate faster and possibly more precise clinical diagnoses and guide treatment strategies in infectious diseases. A medical condition where a faster and more precise diagnosis could make a difference in clinical outcomes is endophthalmitis. Endophthalmitis is an acute intraocular inflammation that can lead to a permanent loss of vision. It often develops in response to microorganisms (usually bacteria and fungi) that enter the eye following eye surgery such as cataract surgery and intravitreal injection. The treatment strategy as well as visual outcome depends in part on the identity of the causative agents. For example, endophthalmitis cases involving coagulase-negative staphylococci have a better prognosis than cases involving enterococci or streptococci ^1^. Often, the involving bacteria appear to originate from the patients’ own microbiota, but may also be introduced through contaminated solutions or instruments used during eye surgery ^2,3^ Endophthalmitis is an acute emergency and therefore clinicians start with a treatment before obtaining information about the identity of the causing microbial agent. It is anticipated that in the future, a more rapid determination of the identity of the causing agents and their antimicrobial resistance profiles using diagnostic metagenomics could facilitate the application of more precise treatments and reduce blindness.

Cataract is a condition in which the lens of the eye becomes progressively opaque and is one of the major causes of reversible visual loss. It is estimated that every year 10 million cataract surgeries are performed around the world ^4^ The risk of endophthalmitis after cataract surgery is 1.4-4 per 10,000 cataract surgeries in the US and Denmark, and can be higher in other countries ^1,5,6^. About 1/3 of the eyes with endophthalmitis in cataract patients remain blind after treatment ^7^

Intravitreal injection with anti-vascular endothelial growth factor (anti-VEGF) has revolutionized the treatment of wet age-related macular degeneration, as well as diabetic maculopathy, and retinal vein occlusions during the last decade. It is the fastest growing procedure in ophthalmology and it was estimated that the number of intravitreal injections in the US would reach nearly 6 million in 2016 ^8^. The risk of endophthalmitis after intravitreal injection is approximately 4.9 per 10,000 intravitreal injections ^9^.

The diagnosis and treatment of endophthalmitis is performed by vitrectomy surgery or a vitreous tap ^10^. A vitrectomy is a procedure in which the vitreous body of the eye, which is the immobile gel-like fluid that occupies the space between the lens and retina, is aspirated and replaced by balanced salt solution. A vitreous tap is a more simple procedure where the vitreous is aspirated without being replaced by balanced salt solution. In both procedures, antibiotics, such as vancomycin combined with ceftazidime, are being injected into the vitreous body to treat the infection. The vitreous is often examined for infectious agents in the clinical laboratory using cultivation-based techniques.

In the clinical setting it is challenging to distinguish between infectious endophthalmitis and non-infectious (“sterile”) endophthalmitis. Studies have shown that the proportion of culture-positive cases can be as low as 39% after cataract surgery and 52% after intravitreal injection ^9,11^. Polymerase chain reaction can increase the rate of identifying the microorganisms by 20% ^12^, but in many endophthalmitis cases a causative agent cannot be identified. It is also unclear, whether the vitreous in endophthalmitis may contain multiple microorganisms that are not all being detected with the current methods. Furthermore, from a clinical perspective it is of importance to have a method that facilitates the identification of the cases of non-infectious endophthalmitis. Non-infectious endophthalmitis can present as a variant of TASS (toxic anterior segment syndrome), and these patients may benefit from steroid instead of antibiotic treatment to obtain a better visual outcome ^13^.

Genomics approaches have the potential to revolutionize clinical diagnostic and therapeutic approaches in particular in the area of infectious diseases. Using shotgun metagenomic sequencing, a range of microorganisms and possible causing agents (e.g. bacteria, archaea, fungi, protozoa, viruses) can be identified ^14,15^. In addition, upon cultivation-based isolation of microorganisms from the patient specimen, these can be subjected to whole genome sequencing (WGS) and *in silico-*determination of their taxonomic affiliation, phylogenetic relationships, potential antibiotic resistance genes, and virulence-associated genes ^16,17^

Here, we perform metagenomic sequencing of vitreous specimens obtained from patients with endophthalmitis and a range of control samples. We evaluate two DNA isolation procedures for vitreous, and describe a bioinformatics workflow for data analysis and identification of potential infectious agents. The workflow includes *in silico* filtering steps for the removal of human DNA sequences, ambiguous and contaminant sequences in reference genomes from public repositories, and background DNA detected in control samples. We compare the metagenomics-based results with the results from the routine clinical cultivation-and mass spectrometry-based analysis, as well as to WGS-based identification of isolates obtained from the vitreous. Our findings suggest that metagenomics analysis together with WGS-based analysis is suitable for the identification of the potential infectious agents from human ocular body fluid, and in the future could guide therapeutic strategies including targeted antimicrobial therapy and the choice of steroids.

## Results

### Study design and metagenomic sequencing

To evaluate the use of shotgun metagenomic sequencing for the identification of potential disease-causing agents in postoperative endophthalmitis, we collected vitreous during vitrectomy from 14 patients with endophthalmitis (7 post cataract surgery, 7 post intravitreal injection) (Figure 1, Supplementary Table S1). As control, we obtained vitreous from 7 patients without endophthalmitis during macula hole surgery. Additional controls included 6 balanced salt solution (BSS) aliquots, of which 3 originated from individual bottles (BSS-B), and 3 from the vitrectomy BSS infusion lines (to be inserted into the eye) after the bottle had been connected to the vitrectomy system (BSS-S) (Figure 1). As there exist no standard procedure for the isolation of DNA from vitreous, we examined two procedures using the QIAamp DNA Mini Kit (QIA) and QIAamp UCP Pathogen Mini kit (UCP), and 4 extraction (blank) controls were included per kit (Figure 1).

**Figure 1:**
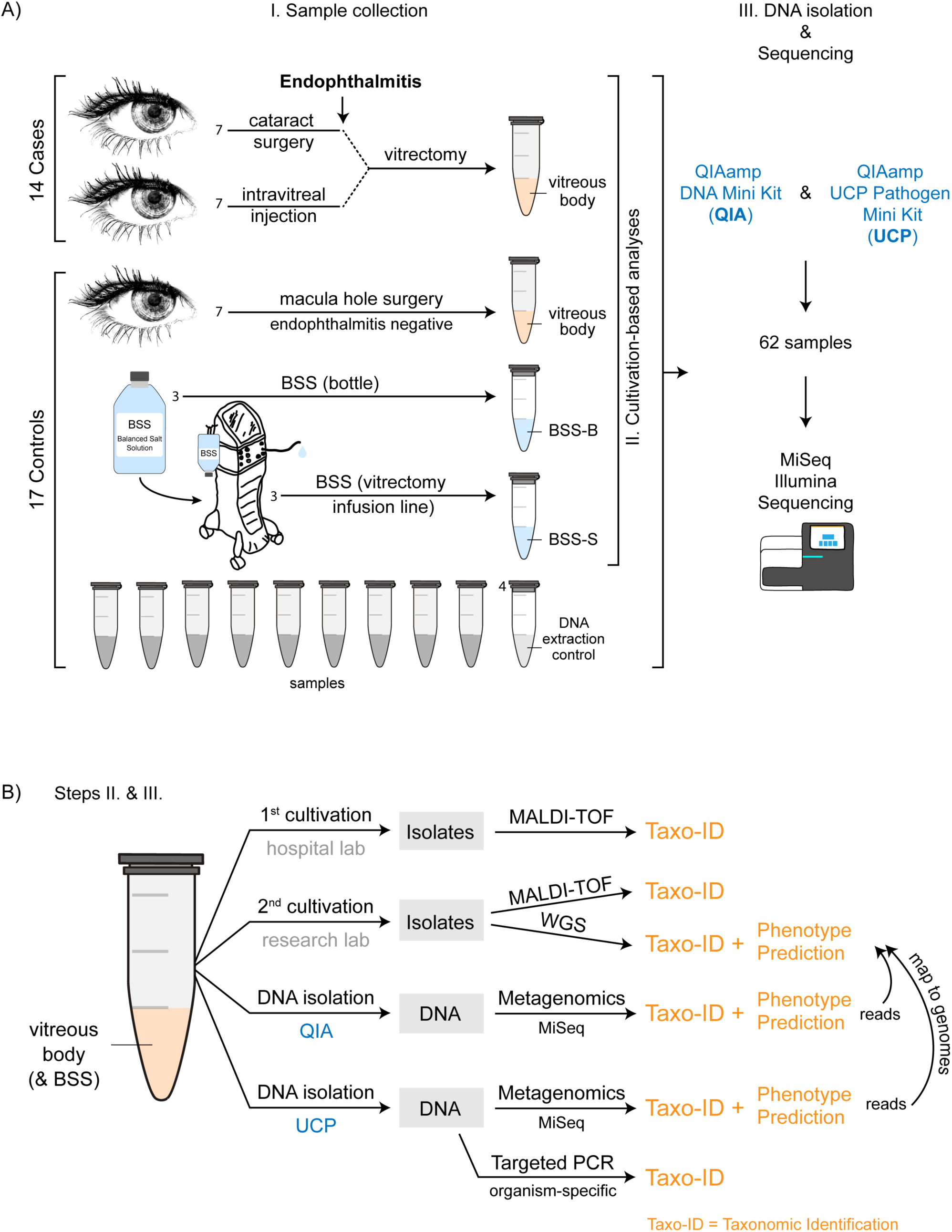
Sample collection, DNA isolation, and shotgun metagenomic sequencing. A) I.) Sample collection: Vitreous body (intraocular body fluid) was collected through vitrectomy from 14 patients with endophthalmitis following cataract surgery (n=7) and intravitreal injection (n=7). As control, vitreous was collected from 7 patients without postoperative endophthalmitis during macula hole surgery. Six aliquots (3 sample pairs) were obtained from balanced salt solution (BSS) that is infused into the eye during vitrectomy. Three aliquots were collected from separate BSS bottles (BSS-B), and the second set of aliquots was collected from the vitrectomy surgical system (BSS-S) after it had passed through the vitrectomy infusion line, respectively. The samples were examined using II.) Cultivation-based analyses and III.) DNA isolation (2 methods) & Metagenomic shotgun sequencing, including the examination of DNA extraction (blank) controls. A total of 62 samples were sequenced using Illumina MiSeq sequencing technology. B) More details to steps II.) and III.): II.) Cultivation-based analyses: Aliquots of the vitreous body fluid and balanced salt solution samples were subjected to cultivation-based analyses separately at the hospital and research laboratories. Obtained isolates were analyzed using mass spectrometry and whole genome sequencing. III.) DNA isolation & Metagenomic shotgun sequencing: Samples were extracted using two DNA isolation procedures: QIAamp DNA Mini Kit (QIA), and QIAamp UCP Pathogen Mini kit (UCP). A DNA extraction (blank) control was included at each round of DNA isolation, i.e. one DNA extraction control for 12-14 samples in total per extraction round (more vitreous samples were extracted than analyzed in this study). To verify the presence of the main microorganisms detected in the metagenomics analysis, the shotgun metagenomics reads were mapped to the genome assemblies of the isolates obtained from the vitreous samples. Not displayed here is the mapping of metagenomic shotgun reads to microbial reference genomes in the database (Provided in Figure 4). As an additional verification, PCR analyses were carried out to detect the presence of the most abundant microorganisms in the vitreous samples using organism-specific primer sets.

The 62 samples were sequenced using Illumina MiSeq sequencing technology and a total of 90.6 million raw read-pairs were obtained. The average number of read-pairs after quality control for the endophthalmitis patients were 2.1/2.3 million read-pairs (QIA/UCP), and for the endophthalmitis-negative vitreous samples 1.0/0.6 million read-pairs (QIA/UCP). The average number of read-pairs for the BSS samples were 52,899/6,067 (QIA/UCP), and for the DNA extraction controls 20,931/3,134 (QIA/UCP). Overall, more read-pairs were obtained on average for the control samples when extracted with the QIA kit, while more read-pairs were obtained for the vitreous from the endophthalmitis patients when extracted with the UCP kit (Supplementary Fig. S1, Supplementary Table S2).

### Identification of human-affiliated DNA sequences

In a first-pass analysis, in which we mapped the reads against a set of reference genomes, we detected a high number of reads affiliated with human DNA sequences, which was anticipated in particular in the endophthalmitis cases that can experience an infiltration of immune cells into the vitreous chamber. Hence, we implemented a 2-step filtering process to remove the reads affiliated with human genome sequences (Figure 2). In the first step we removed the reads that mapped to the human reference genome (GRCh8.p10). Due to the genetic individuality of humans some reads might not map to this reference genome, and therefore we removed in a second step all reads that aligned to any human DNA sequence entry in the NCBI nt database (Supplementary Fig. S2, Supplementary Table S2).

**Figure 2:**
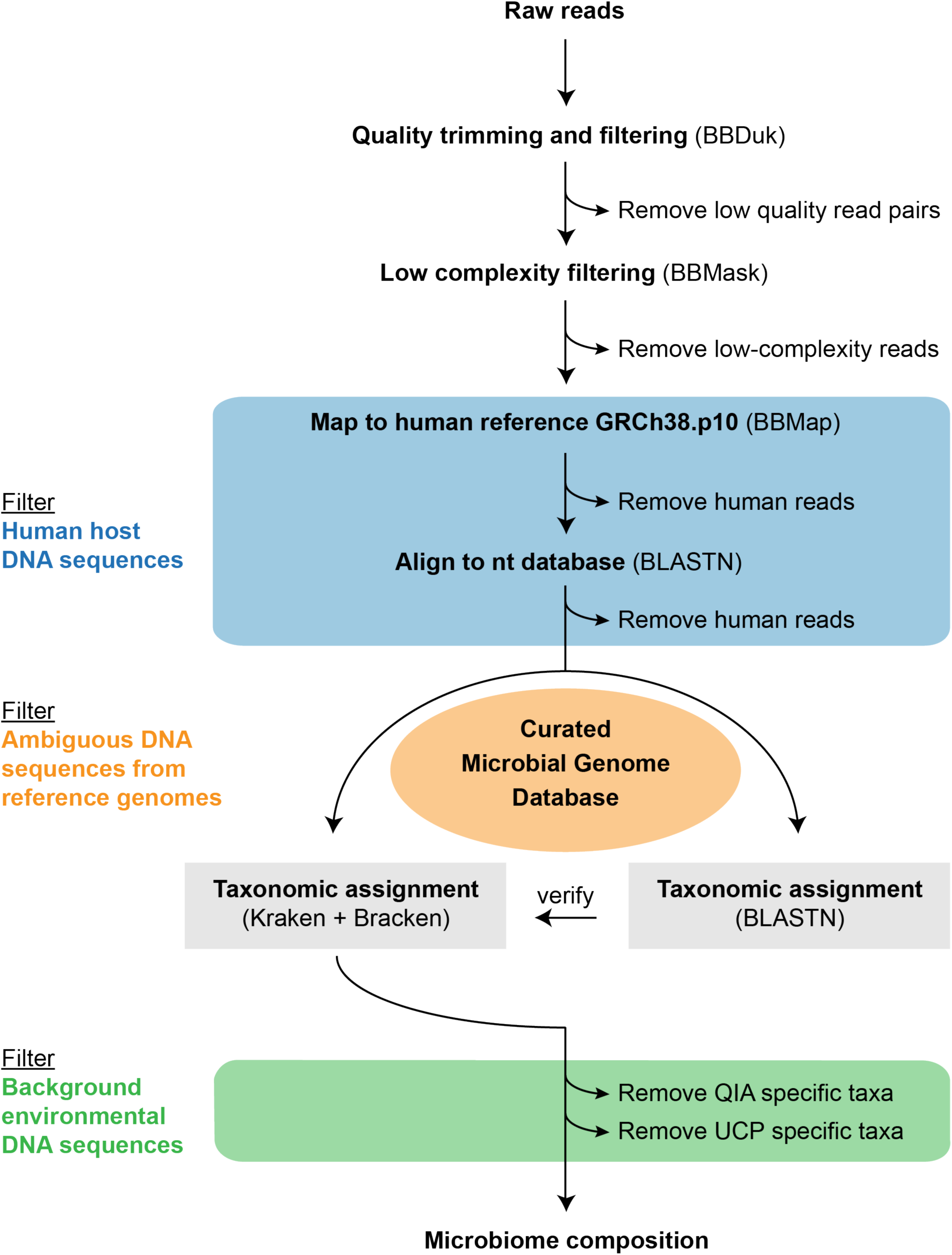
Workflow for metagenomic data analysis. In a first step, sequencing adapters, low quality bases, and reads with low complexity were removed. Subsequently, reads that mapped against the human reference genome sequence, or aligned with human sequences in the nt database were removed. The taxonomic classification of the reads was performed with Kraken together with Bracken using a curated microbial genome database containing 5750 microbial (archaea [251], bacteria [5166], fungi [225], protozoa [73], viruses [35]) and 1 human reference genome sequence (for details, see Supplementary Methods). Additional reads that in this step were classified as human were removed. To verify the classification results, the reads were also aligned to the reference genomes using BLASTn. Organisms specific for the DNA extraction (blank) controls were filtered from the patient samples.

### Identification of ambiguous and contaminant DNA sequences in genomes from public repositories

In the initial first-pass analysis involving mapping of reads against reference genomes, we observed that some genomes recruited particular high numbers of reads. These included *Hammondia hammondi* strain H.H.34, *Alcanivorax hongdengensis* Strain A-11-3, *Toxoplasma gondii* ME49, and *Arthrobacter* sp. Soil736. Upon inspection of these genomes we found that the reads mapped only to specific genome sequence fragments such as short contigs and scaffolds (Supplementary Fig. S3). To examine why specific contigs and scaffolds recruited high numbers of reads, we aligned these against the nucleotide collection nt (NCBI). We found that the Top10 matches for most of these contigs and scaffolds included several human DNA sequence entries that are not part of the human reference genome GRCh8.p10 (Supplementary Table S3). While a few scaffolds of *Hammondia hammondi* strain H.H.34 aligned with human DNA sequences (e.g. scaffold NW_008644893.1), many aligned to *Bradyrhizobium* spp. genomes in the nt database (Supplementary Table S3), indicating that human as well as microbial sequence contamination can be found in public genome assemblies.

### Construction of a curated microbial genome database

Our analysis suggested that some microbial reference genomes contain ambiguous/contaminant sequences and we aimed at constructing a curated microbial genome database, devoid of these sequences to the extent possible. Removing these sequences could reduce the number of false positive hits that are the result of either contaminant sequences in the (incomplete) genome assemblies, or because highly similar sequence regions naturally exist across genera that result in the classification of reads to a different genus. We examined 5715 of the microbial reference and representative genomes (archaea, bacteria, fungi, protozoa) (Supplementary Table S4) and aligned all sequences ≤10 kb against the nucleotide collection nt (for a detailed description, see Supplementary Methods). A total of 70,478 ambiguous sequences (contigs and scaffolds) were identified, of which the majority were detected in incomplete microbial genomes. A total of 62% of all incomplete microbial genomes had sequences flagged as ambiguous (range: 1 - 10,590; average: 28 sequence fragments). Ambiguous sequences were identified in 43% of all bacterial and 72% of all protozoan genomes, and on average comprised 0.36% and 0.84% of the total genome sequence, respectively (Table 1, and https://figshare.com/s/a282670f1405eae232df, https://figshare.com/s/045b1252bd7555b50ef0, https://figshare.com/s/c42158cdee23f25489cd) ^18^. The ambiguous sequences were removed and the resulting reference microbial genome database contained a total of 5,751 genomes with 34 Tb (including 3.1 Tb for the human genome). The code for the creation of the curated microbial reference genome database is accessible from Github (https://github.com/philDTU/endoPublication), and the curated microbial reference genomes can be downloaded from ftp://ftp.cbs.dtu.dk/public//CGE/databases/CuratedGenomes.

**Table 1.** Ambiguous, including contaminant, sequences in public microbial genomes.

|  | Total |  | Ambiguous sequences* |  |  |
| --- | --- | --- | --- | --- | --- |
|  | Genomes | Bases | Genomes | Bases | Bases (%) |
| Archaea | 251 | 673,145,451 | 65 | 1,813,095 | 0.27 |
| Bacteria | 5166 | 20,854,687,300 | 2251 | 75,888,994 | 0.36 |
| Fungi | 225 | 6,486,874,847 | 126 | 6,642,500 | 0.10 |
| Protozoa | 73 | 2,930,167,033 | 53 | 26,447,579 | 0.84 |
| Viruses | 35 | 640,331 | ND | ND | ND |
| Sum | 5750 | 30,945,514,962 | 2922 | 110,858,157 | 0.35 |

\*Genomic sequence regions ≤10 kb (incl. contigs and scaffolds) that had a match (e-value ≤ 1e-6; query coverage ≥ 70%) belonging to a different genus than their stated genus definition when aligned against the non-redundant nucleotide collection (nt) database from NCBI. For more details, see Supplementary Methods and https://figshare.com/s/a282670f1405eae232df, https://figshare.com/s/045b1252bd7555b50ef0, https://figshare.com/s/c42158cdee23f25489cd. ND=not determined

### Identification of contaminant (environmental background) DNA sequences in samples

From the sequencing of DNA extraction (blank) control samples we obtained sequencing data, albeit at a lower frequency compared to the patient specimens (Supplementary Fig. S1). The *in silico* identification and removal of background DNA sequences are of critical importance, particularly from specimens where the potential infectious agent may be present in low abundance. We carefully examined the eight DNA extraction control samples and devise a list of the most abundant and frequent environmental contaminant taxa in these samples (Supplementary Table S5, Supplementary Fig. S4). We did not include taxa in the list that were occasionally observed in endophthalmitis-positive patients and that were detected at a higher abundance in these samples than in the respective DNA extraction controls. These non-contaminant taxa include *Enterococcus faecalis, Escherichia coli, Micrococcus luteus, Staphylococcus aureus*, and *Staphylococcus epidermidis* (Figure 3, and https://figshare.com/s/a4fd9d84260e8456ab72). The microbial composition patterns in the DNA extraction control samples appeared to be influenced by the choice of DNA isolation kit, the day of DNA extraction, and sequencing run (Supplementary Fig. S4). The contaminant taxa (Supplementary Table S5) were removed from the datasets of all endophthalmitis patients.

**Figure 3:**
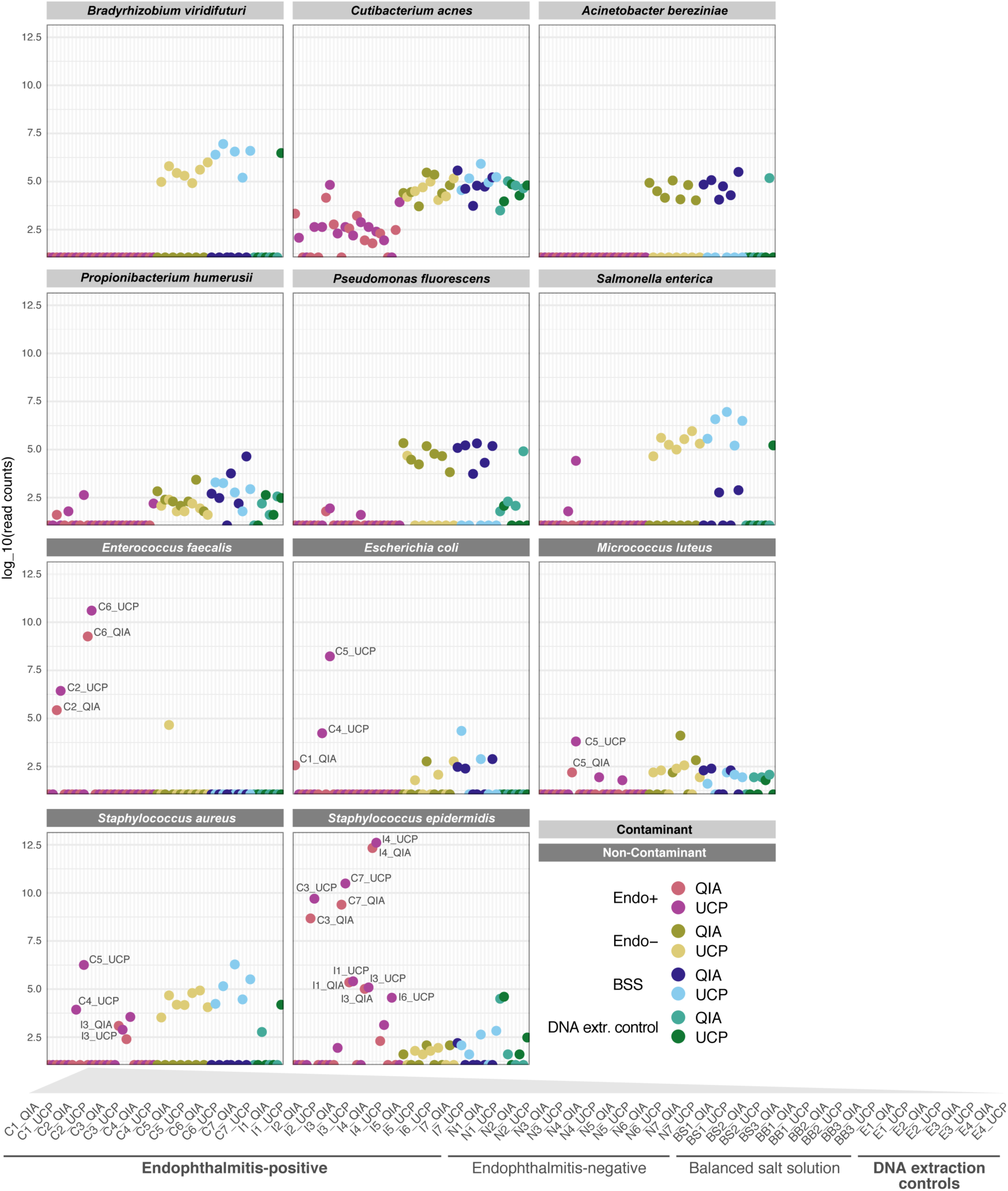
Selected contaminant and non-contaminant organisms based on evaluation of DNA extraction control samples. Contaminant organisms (light grey) were present in higher abundance in DNA extraction controls (green) compared to the endophthalmitis-positive samples (red). The contaminant organisms were detected in similar abundance in the endophthalmitis-negative (yellow) and/or balanced salt solution samples (blue) as in the DNA extraction control samples. Organisms that were detected in higher abundance in patient samples (dark grey), compared to their respective DNA extraction control samples, were not regarded as sample contaminants. Read counts are presented as counts per million in relation to the total non-human read counts per sample, respectively. An interactive version of this figure that includes individual sample information, including read counts, is available from https://figshare.com/s/a4fd9d84260e8456ab72. For a detailed list of contaminant organisms, see Supplementary Table S5.

### The microbial composition in endophthalmitis-negative and balanced salt solution samples is similar to DNA extraction controls

The contaminant taxa that were identified in the DNA extraction controls were often present at similar abundances in the endophthalmitis-negative (vitreous control) and balanced salt solution samples (Figure 3, Supplementary Fig. S5). We found certain taxa to be specific for the DNA isolation method (QIA or UCP) in round C of DNA extractions (Supplementary Fig. S5, Supplementary Table S2). Samples processed using the QIA method contained *Pseudomonas* spp., *Acinetobacter* spp., and *Janthinobacterium* spp. among others, and samples processed with the UCP method included mainly *Bradyrhizobium* spp. Other organisms appeared to be present across all samples (Supplementary Fig. S5). For example, *Cutibacterium acnes* and *Propionibacterium humerusii* were detected in most samples and they might represent environmental bacteria originating from the staff handling the samples or fomites such as the laboratory equipment and supplies.

### Microorganisms in endophthalmitis-positive patients as determined by metagenomics

For 12 out of 14 endophthalmitis patients a dominant microorganism was identified in the vitreous (for all UCP-extracted, and most QIA-extracted specimens) using the read classification approach (Figure 4 and 5). These organisms included *Staphylococcus epidermidis* (six patients), *Enterococcus faecalis* (two patients), *Serratia marcescens* (one patient), *Paenibacillus* spp. (one patient), and *Staphylococcus hominis* (one patient). In one patient (C5), a number of different organisms were identified, most dominantly *E. coli* in the UCP-extracted specimen (>3000 reads), *Moraxella catarrhalis* (11 reads) in the QIA-extracted specimen, and *Micrococcus luteus* with 9 and 45 reads in QIA and UCP-extracted samples, respectively (Figure 4 and 5, and https://figshare.com/s/5feabfad1d8c495bf7a3). For two additional patients, *Commamonas testosteronii* and *Escherichia coli*, or *Caulobacter* spp. were identified as the most dominant organisms respectively (C1, I7), however, these were only represented by <25 reads. In the seven patients that contracted endophthalmitis following cataract surgery, the most frequent bacteria were *Enterococcus faecalis* (two patients), *Staphylococcus epidermidis* (two patients), and *Serratia marcescens* (one patient). In the seven patients with endophthalmitis following intravitreal injection, the most frequent bacteria were *Staphylococcus epidermidis* (four patients), *Paenibacillus* spp. (one patient), and *Staphylococcus hominis* (one patient) (Figure 5). Overall, potential causing agents where identified with 58 reads (*Paenibacillus* spp.) as a lower bound in patient I2, and 2,999,838 reads as the highest detected read count (*Staphylococcus epidermidis*) in patient I4. The presence of the two most frequently detected bacteria, *Staphylococcus epidermidis* and *Enterococcus faecalis*, in the vitreous fluid was also verified using targeted PCR assays (See Supplementary Methods, and https://figshare.com/s/0e8a98f436f07efc4dd5).

**Figure 4:**
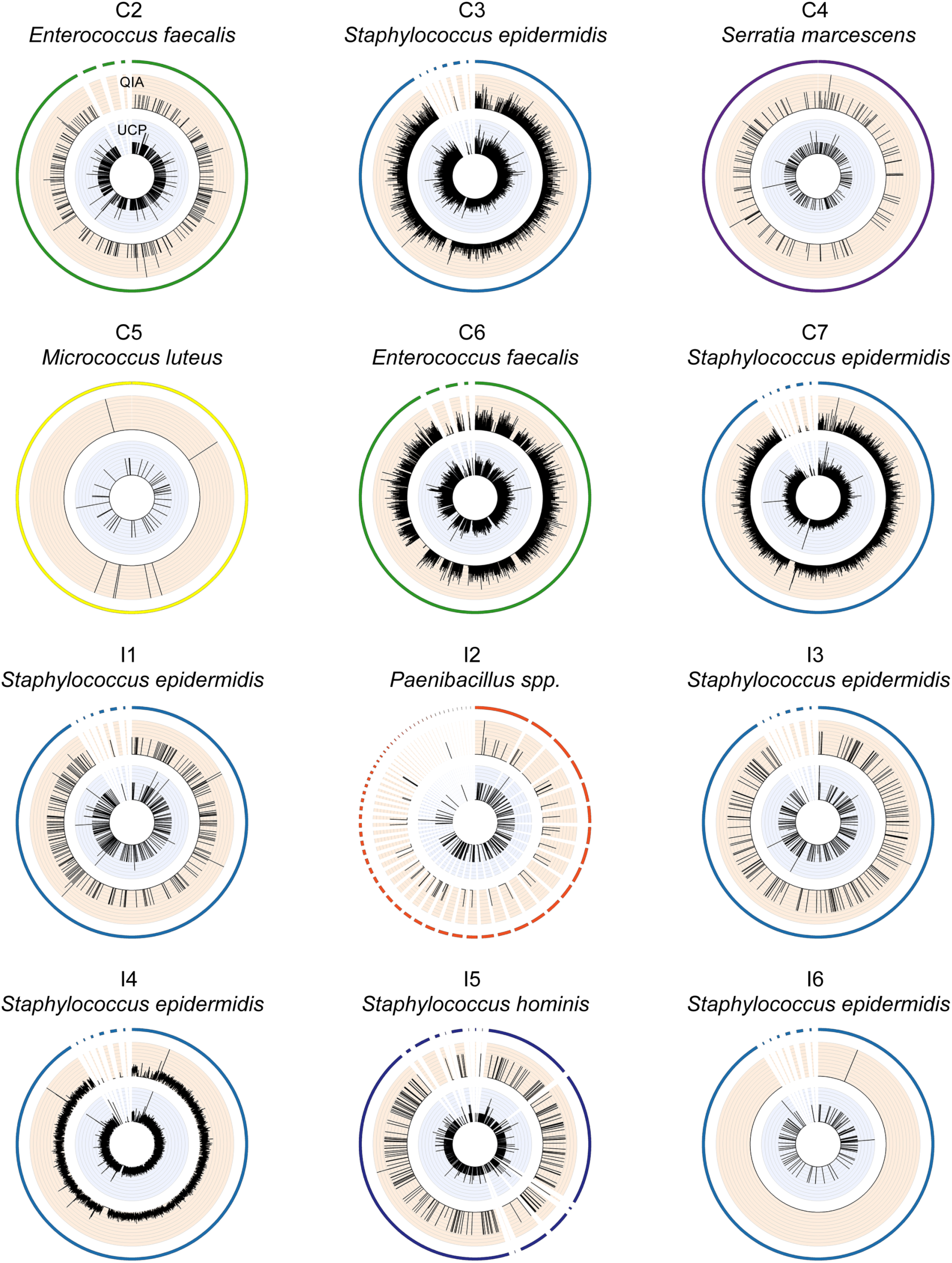
Coverage of bacterial reference genomes by metagenomic reads originating from intraocular fluid of endophthalmitis patients. For each individual patient, the metagenomic shotgun reads of the most abundant microbial organism were extracted at genus-level and mapped as unpaired reads using BBmap suite to the respective reference genome sequence in the database. For patients C1 and I7 a particular microbial organism could not be assigned confidently in the metagenomics analysis, and these patients are regarded as “sterile” endophthalmitis cases. The outer most circle displays all sequences of the reference genome (including short contigs and plasmids). The orange and blue inner circles display the depth of mapped reads originating from the vitreous specimens that were extracted with the QIA and UCP DNA extraction methods, respectively. In the two cases where metagenomics analysis and culture results from the hospital were not identical regarding the most abundant organism (patients C5 and I3), we examined the reads via genome mapping for all organisms detected in the metagenomics analysis. The most relevant abundant organism is shown here and the additional plots, as well as information about the maximum read depth for all detected organisms, is available from https://figshare.com/s/c2ce2d32daf25db54904.

**Figure 5:**
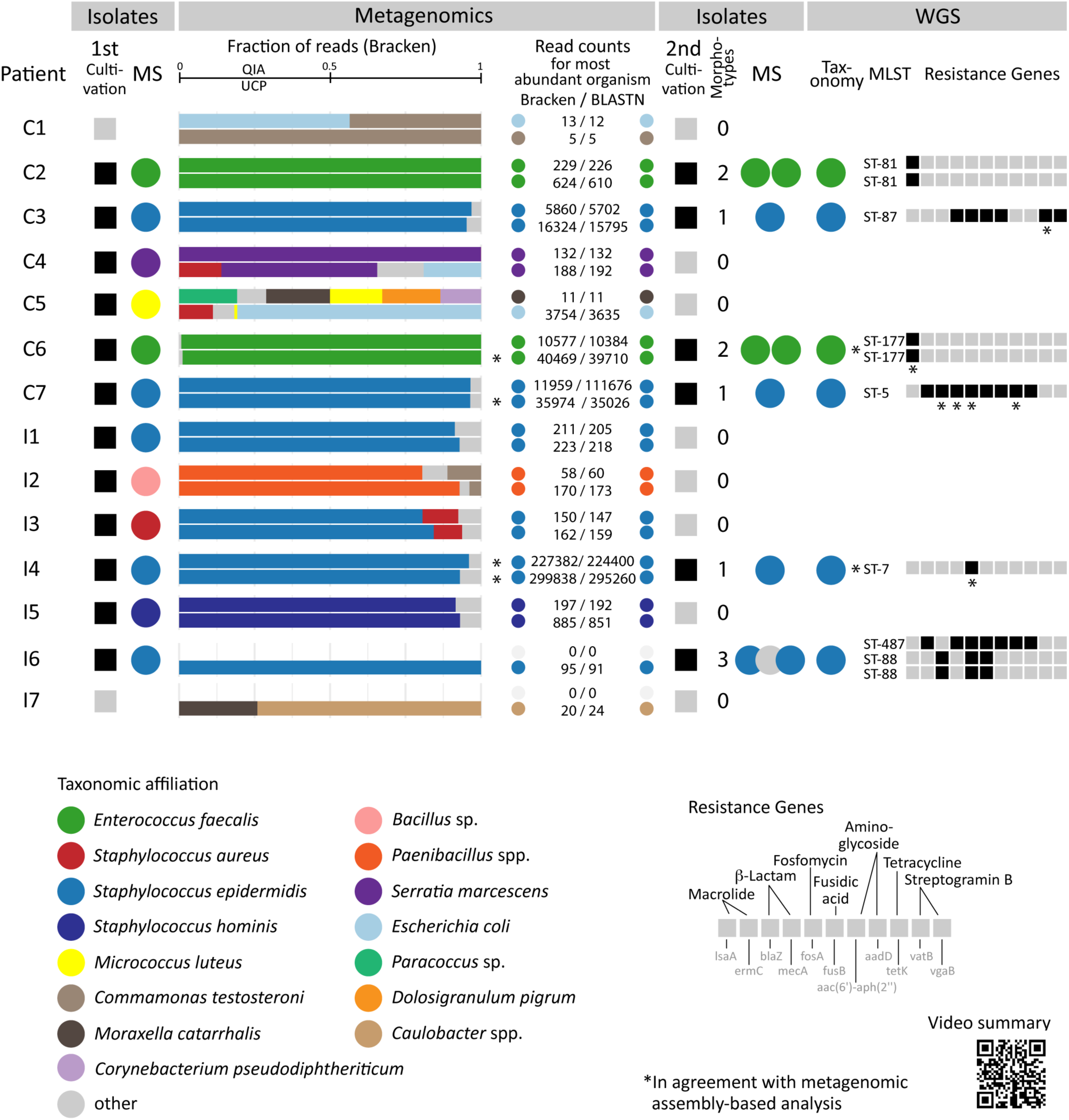
Summary of cultivation-based, metagenomics, and whole genome sequence analyses. Bacterial isolates were obtained at the hospital laboratory (1^st^ cultivation) from vitreous from endophthalmitis patients following cataract surgery (C1-7) and intravitreal injection (I1-7) and the taxonomic affiliation of the isolates were determined by MALDI-TOF mass spectrometry (MS). Vitreous was analyzed through metagenomics at the research laboratory using two DNA isolation methods (QIAamp DNA Mini Kit, QIA; QIAamp UCP Pathogen Mini kit, UCP) and the taxonomic affiliation of reads was determined. The detected amount of human DNA sequences in percent (%) is provided in the first column of the Metagenomics tab. In the horizontal bar charts, the taxonomic identity and relative fraction of microbial reads for the most abundant identified organisms based on the Kraken+Bracken analysis is indicated for both DNA isolation methods. The read counts for the most abundant organism according to the Kraken+Bracken (all reads) and BLASTN (forward read) analyses are indicated to the right. The read counts for the most abundant organisms per sample as determined by Kraken, Bracken, and BLASTn analyses are available through figshare at https://figshare.com/s75feabfad1d8c495bf7a3. Bacterial isolates for some samples were obtained in a second round of cultivation at the research laboratory (2^nd^ cultivation), and one representative per colony morphotype per vitreous sample was subjected to MS and whole genome sequencing (WGS). The taxonomic affiliation of isolates was determined through classification of assembled genomes using a k-mer based approach and genomic MLST, and antibiotic resistance genes were identified using ResFinder. Furthermore, metagenomic assemblies were generated from the shotgun metagenomic reads and analyzed with regards to taxonomic affiliation and selected functional characteristics (Supplementary Table S6). A video summary is available from figshare at https://figshare.com/s/38fe043f6a8ef1710444.

In addition to the read classification approach, we constructed metagenomic assemblies for the individual samples and characterized these according to a number of taxonomic and functional categories, including bacterial species affiliation, sequence type, genomic MLST, resistance genes, virulence-associated genes, and plasmids. For the three patients for whom we obtained high numbers of classified reads using the read classification approach described above (C6, C7, I4) (Figure 5), we obtained information in nearly all categories using the metagenomics assembly approach (Supplementary Table S6). The taxonomic information that we obtained using the metagenomic assembly approach was in agreement in all cases with the taxonomic information we obtained using the metagenomic read classification approach. Furthermore, using metagenomics assembly analysis we detected a number of antimicrobial resistance genes in the specimens for which we also obtained taxonomic information. In addition, we detected a streptogramin B resistance gene (vat(B)) in sample C3_UCP, an aminoglycoside resistance gene (aadD) in sample C7_QIA, and a Col plasmid origin of replication in sample C5_UCP. Of note, five out of the seven total samples, for which we obtained information using the metagenomics assembly approach, were ocular body fluid samples that had been processed using the UCP DNA isolation protocol (Supplementary Table S6).

### Bacterial isolates from endophthalmitis patients have in most cases the same identity as the most abundant organism determined by metagenomics

At the hospital microbiology laboratory, bacteria could be isolated from the vitreous right after vitrectomy for 12 out of 14 patients. The identity of the isolates was determined by MALDI-TOF mass spectrometry (MS), and in nine cases the same agent was identified as in the metagenomic analysis (Figure 5). In addition, *Micrococcus luteus* was isolated from patient C5 in both QIA‐ and UCP-extracted samples (9/45), but this organism was not the most abundant one identified using the metagenomics sequencing-based method (Figure 5). Using the cultivation-based method a *Bacillus* sp. (Order: Bacillales) was determined for patient I2, and reads classified as *Paenibacillus* spp. (Order: Bacillales) were identified using the metagenomics analysis. A *Staphylococcus aureus* culture was obtained in the hospital for patient I3, and *S. aureus* was also represented with 22/18 (QIA/UCP) reads in the metagenomics analysis in this patient, even though *S. epidermidis* was the most abundant organisms identified using this approach (150(QIA)/162(UCP) reads). In the two cases for which the culture-based approach was negative (C1, I7), only fewer than 25 reads were classified using the metagenomics approach.

At the research laboratory, we attempted a recultivation of microorganisms from frozen vitreous and successfully obtained isolates for six patients. Different colony morphotypes on the agar plates were obtained and analysed using whole genome sequencing (WGS) and MALDI-TOF mass spectrometry (MS). These isolates had the same species affiliation according to WGS and MS analyses as the isolates obtained at the hospital for the same patient, and as identified in the vitreous using metagenomics analysis (Figure 5). The presence of the isolates in the vitreous samples was further verified by mapping the shotgun metagenomic reads originating from the vitreous to the genomes of the isolates, and an even breath of coverage was observed for all isolates (https://figshare.com/s/c2ce2d32daf25db54904).

Using WGS we found that for multiple morphotypes the same organism and sequence type was identified, with one exception. For patient I6, we obtained three *Staphylococcus epidermidis* isolates and of which two belonged to sequence type ST-88 and one to ST-487. The *Staphylococcus epidermidis* isolates obtained from other patients (C3, C7, I4) belonged to different sequence types (Figure 5, Supplementary Table S7), suggesting that they have different origins. Each *Staphylococcus epidermidis* sequence type exhibited its own set of antibiotic resistance genes, including genes facilitating resistance to macrolides, β-lactams, aminoglycosides, and tetracyclines. The *Enterococcus faecalis* from two patients (C2, C6) belonged to different sequence types, and both sequence types shared a gene facilitating resistance towards macrolides (Figure 5, Supplementary Table S7). Several resistance genes that were identified in the sequenced isolates were also identified in the metagenomic assembly analysis (Figure 5, Supplementary Table S6). Some of the resistance genes and their predicted functions identified using the genomics approaches were also in alignment with results from the phenotypic antibiotic susceptibility testing of the isolates obtained during the 1^st^ culturing at the hospital (https://figshare.com/s/e579abea97dfc8c77a6a).

### Detection of bacteriophages and human DNA viruses

As we did not identify a dominant microorganism in two endophthalmitis patients (C1, I7) we examined whether these or any of the other specimens contained DNA viruses not represented in our microbial genome database. We added an additional 7,180 virus genome sequences to the 35 RefSeq virus genomes (https://figshare.com/s/b040289827b79d3a60df) in our database and classified our metagenomic sequencing data using kraken. We identified several *Enterococcus*, *Staphylococcus* and *Propionibacterium* bacteriophages among others in specimens that also were identified to contain the respective bacterial host (https://figshare.com/s/ff0527509828d1529ad9). To evaluate whether our metagenomics approach (Figure 2) would facilitate the identification of human DNA viruses, we analysed metagenomic data obtained from patients with uveitis in which human DNA viruses had been detected previously ^19^. We obtained similar results as described by Doan and colleagues, including the identification of herpes simplex virus 1 (HSV-1) in subject 1, and rubella virus in subject 6 (Supplementary Table S8). In the previous study *Hammondia hammondi* was identified in subject 3 as the second most abundant organism after *Toxoplasma gondii*. We also detected *Toxoplasma gondii* as the most abundant organism in this specimen (represented by 4,410 reads), but did only detect 4 reads for *Hammondia hammondi*; most likely because we had removed DNA sequences from the *Hammondia hammondi* genome in the database that were flagged as ambiguous. In addition, in subject 5 we detected *Ochrobactrum anthropi*, an agent that had been identified previously in eye infections such as endophthalmitis and keratitis ^20,21^. However, we detected *Ochrobactrum anthropi* in high abundance in the water control sample, and therefore it may here rather represent an environmental contaminant.

## Discussion

Metagenomic sequencing-based analyses of complex patient specimens and whole genome sequencing (WGS) of microbial isolates will advance clinical diagnostics and treatment strategies in infectious diseases ^22-24^. One example, for which this strategy may be advantageous is postoperative endophthalmitis as currently a causing microbial agent can only be identified in a fraction of these cases ^1^. Immediate diagnosis and treatment of endophthalmitis is required to prevent vision loss of the affected eye, and it would be helpful to be able to distinguish between infectious and non-infectious (“sterile”) endophthalmitis. Challenges in clinical metagenomics remain at several levels, from specimen collection and processing to the generation of actionable information. We examine here vitreous samples from endophthalmitis patients together with a variety of control samples, evaluate two DNA isolation procedures, create a curated microbial reference database, and present a workflow for metagenomic sequencing data analysis. We compare the results from metagenomic read analysis to WGS and MALDI-TOF mass spectrometry identification of isolates obtained for several patients, as well as to results from metagenomic assembly analysis.

Vitreous samples were collected from 14 patients with endophthalmitis. Seven patients developed endophthalmitis post cataract surgery, in which the natural intraocular lens was exchanged with an artificial one, without the introduction of surgical instruments into the vitreous body. Another seven patients developed endophthalmitis post intravitreal injection, a procedure in which drugs were introduced into the vitreous body using surgical instruments to treat retinal diseases such as age-related macular degeneration. As controls, we included i) vitreous samples from seven endophthalmitis-negative patients, ii) balanced salt solution used during vitrectomy from both, individual bottles and the vitrectomy system after the solution had passed the vitrectomy infusion lines, and iii) DNA extraction (blank) controls (Figure 1).

We investigated two DNA isolation procedures, QIAamp DNA Mini Kit (QIA) and QIAamp UCP Pathogen Mini kit (UCP), for metagenomics analysis to determine possible infectious agents in the vitreous fluid. We obtained more reads (total and classified) on average for endophthalmitis-positive specimens when vitreous fluid was extracted with the UCP kit compared to the QIA procedure (Supplementary Figure S1 and Figure 5). In contrast, lower numbers of reads on average were obtained from the three types of control samples when they were extracted with the UCP kit compared to the QIA procedure (Supplementary Figure S1). Our analysis revealed that UCP-extracted control samples harboured a lower microbial diversity compared to QIA-extracted ones (Supplementary Figures S4 and S5). Even though we identified distinct QIA and UCP kit “fingerprints”, bacteria such as *Cutibacterium acnes* and *Propionibacterium humerusii* were present as background DNA across samples, independent of the DNA isolation kit. These bacteria likely originated from the staff handling the samples and/or additional laboratory supplies that were used during sample handling and processing ^25^. Contaminant background DNA has been identified previously in other DNA isolation kits ^26-28^, and our analysis suggests that an ultraclean production of reagents and consumables reduced the amount of background DNA in the UCP DNA isolation kit reagents and/or supplies. Contaminant viral DNA has been identified in previous sequencing-based studies as well such as hybrid parvovirus-like virus NIH-CQV/PHV DNA from silica column-based nucleic acid isolation kits ^29,30^, and which we detected in our samples, too (https://figshare.com/s/ff0527509828d1529ad9). We did not, however, detect torque teno virus DNA, as previously described for some endophthalmitis cases ^31^. Overall, the UCP kit appeared to be suited for the isolation of DNA from vitreous, and may potentially be useful for other human body fluids and biological specimens that are assumed to have a low microbial biomass.

Our metagenomics data analysis workflow included three filtering steps (Figure 2) to reduce i) human host DNA sequences, ii) false positive hits due to ambiguous and contaminant DNA sequences in reference genomes, and iii) environmental background DNA sequences introduced by kit reagents, potentially other laboratory supplies, as well as laboratory staff. We particularly found that ambiguous/contaminant sequences in public genomes, which serve as reference in many metagenomic studies, could lead to the false positive identification of microorganisms. Our initial read classification, in which we used the original reference genomes, revealed *Toxoplasma gondii* (false positive) across samples, even after filtering reads that mapped to the human reference genome. Some microbial reference genomes appeared to harbour human DNA sequences not present in the human reference genome, thus making it challenging to detect these sequences in the initial human DNA sequence filtering step. This effect is especially critical when analysing clinical specimens, since the patient’s DNA is expected to be found in these samples. In addition, we noticed that certain microbial genomes contained sequences that had a high similarity to other microorganisms belonging to a different genus. These can be correct naturally occurring DNA sequence regions that have a high similarity across a range of microbial taxa (including regions acquired via horizontal gene transfer). In other cases they can be contaminant contigs or scaffolds in primarily incomplete genome sequence assemblies. In either case, the read classification can lead to a false positive identification of microorganisms. Contaminant DNA sequences in published genomes have been previously found, particularly in human and animal genome assemblies ^32-34^. Hence, we systematically examined 5,715 microbial reference and representative genomes (archaea, bacteria, fungi, protozoa), and in 62% of all incomplete microbial genomes sequences were flagged as ambiguous. We removed 70,478 ambiguous sequences (including human and microbial contaminants), reflecting 0.35% of the bases in total from the microbial reference genomes (https://figshare.com/s/045b1252bd7555b50ef0). The majority of the removed sequence fragments are a correct part of the respective genome. However, a more complex and thorough analysis is required in the future to decide whether a particular part should be removed or not. In our case, many of the remove sequences are plasmids or sequence fragments from lower-quality assemblies. Since plasmids are mobile genetic elements it is unknown at what confidence-level they contribute to the taxonomy assignment because some plasmids have a broad host range. The removed sequences from the lower-quality assemblies should be neglectable in our clinical study, since most of the identified infectious agents are not in that category. According to the insight gained from this study we recommend using curated microbial reference genomes in microbiome studies and particularly for the analysis of clinical samples with an assumed low microbial biomass. Additionally, subsequent to filtering we recommend to always check the sequences that were removed. We provide a script that facilitates the generation of curated databases, including the one used in this study (https://github.com/philDTU/endoPublication) as well as the sequences of the curated genomes (ftp://ftp.cbs.dtu.dk/public//CGE/databases/CuratedGenomes).

Our analysis further demonstrated the benefit of including a variety of control samples. In fact, the number of control samples in this study exceeded the number of the main samples under investigation by a factor of 1.2. By analysing vitreous from endophthalmitis-negative patients, aliquots of balanced salt solution (from bottle and vitrectomy infusion line), and blank DNA extraction controls, we determined the background levels of organisms in the respective environments. All control sample types had a similar microbiome pattern, characterized by organisms found in the corresponding DNA extraction (blank) controls (QIA and UCP) and typical skin inhabitants. We did not identify specific microorganisms for the endophthalmitis-negative patients, similar to previous cultivation-based assessments, suggesting that vitreous fluid is a sterile body part or only contains few microbial cells in individuals without eye infections ^35,36^. We also did not identify any specific organisms residing in balanced salt solution that was infused into the patient’s eye, in addition to the ones identified in DNA extraction controls. In all cases, we cannot however exclude that DNA sequences from other microorganisms would have been found if a deeper DNA sequencing had been performed or RNA had been isolated and analysed by deep sequencing. In our analysis of the vitreous fluid from endophthalmitis-positive patients we removed the background contaminant organisms *in silico* that were detected in the respective DNA extraction controls and were not present in higher abundance in the endophthalmitis-positive patients. To trace the origin of detected organisms, including infectious agents, additional controls in future studies could include samples from: i) the patients skin, eye lid, conjunctiva, or other body sites that are in proximity to the surgical site, ii) the surgical instruments, iii) blank tubes and/or devices used for the collection of the patient specimen, as well as iv) reference mock communities with known composition. Careful analysis of control samples may assist in the design of harmonized standards and guidelines for the sequencing-based analysis of clinical samples and other biological specimens.

Through our metagenomics read classification data analysis workflow we identified a single potential causing microorganism in 11 out of 12 culture-positive cases, and which in most cases agreed with the cultivation-based analyses (Figure 5). For patient C5 we did not identify a single potential causing agent and instead obtained different microbiome patterns for the two sequenced aliquots (QIA and UCP). In both samples we detected *Micrococcus luteus*, in alignment with the cultivation-based analysis. However, *Micrococcus luteus* was not the most abundant organism in the metagenomic analysis. *Escherichia coli* was the most abundant organism in the UCP-extracted sample, but which may also be a contamination introduced during sample handling. For patient I2 we revealed *Paenibacillus* spp. as a possible causing agent, whereas in the cultivation-dependent analysis the isolate was identified as a related *Bacillus* sp using MALDI-TOF. For patient I3, our metagenomic analysis suggests a potential infection by *Staphylococcus epidermidis* together with *Staphylococcus aureus*. In the cultivation-based analysis *Staphylococcus aureus* was identified as the potential causing agent. Only a few metagenomic reads were classified for patients C1 and I7, and which were regarded as contaminants. For these two patients no microorganisms could be isolated by cultivation-based methods, neither at the hospital nor the research laboratory. Therefore, these two patients are assumed to have a non-infectious (sterile) endophthalmitis.

Both, the analysis of metagenome assemblies as well as whole genome sequences of isolates can reveal the presence of antibiotic resistance genes that could potentially guide therapeutic treatment strategies, in particular when verified by results from susceptibility analysis of isolates. In addition, the specific sequence type for infectious agents, such as *Enterococcus faecalis*, *Staphylococcus epidermidis*, and other organisms, can be identified and assist in the source tracking and epidemiology of the particular agent. Detailed evolutionary relationships between isolates can be revealed if sufficient genome sequence information has been obtained. Our analysis of the identified *Enterococcus faecalis* and *Staphylococcus epidermidis* suggests that they may originate from the individuals involved in the surgery or the immediate environment, as different bacterial sequence types and resistant profiles were identified across patients. The metagenomics analysis did not reveal these bacteria to be present in the balanced salt solution samples, further pointing towards an acquisition from another source.

Our previous clinical microbiology research on urinary tract infections and diarrhoeal diseases ^37,38^ had some limitations for assessing clinical metagenomics as a technology, and whose analysis could now be improved by new insight gained from this study. For example, the examined urine samples were pre-processed by using a centrifugation step to remove human cells ^37^. In this step, also microbial cells may have been removed that have a similar density than the human cells and/or were attached to these. While this step can be advantageous to limit human contaminant sequences, it could be of interest to examine samples with and without the sedimentation step and using the DNA isolation procedure and/or data analysis pipeline described in the present study. Furthermore, the presence of potential contaminant DNA sequences was not examined in the previous study. In the study concerning diarrhoeal diseases one challenge was to differentiate between natural intestinal inhabitants, possible infectious agents, and potential contaminants ^38^. Careful bioinformatics filtering steps and inclusion of control samples, as used in the present study, might allow for more robust identifications in the future. To facilitate a more standardized workflow for sample analysis, we have created a list of recommendations for the design and execution of metagenomic sequencing projects (https://figshare.com/s/2a0709b1f0c5e18754df), in addition to specific details described in this study.

In summary, we find that metagenomics analysis, supported by WGS of isolates, may be a promising strategy for the identification and characterization of infectious agents from human ocular body fluid. This technology may also facilitate a more robust differentiation between infectious and non-infectious (“sterile”) endophthalmitis. Nucleic acid extraction from patient specimens, followed by high-throughput sequencing may ultimately provide more rapid insight in regard to the identity of the causing agent(s) than cultivation-based techniques, in particular in light of recent developments in long-read nanopore sequencing and real-time analysis ^39-41^. In cases where the metagenomic sequencing depth of coverage of the microorganism is sufficiently high, valuable functional information such as antibiotic resistance and virulence-associated genes can be revealed. Prerequisites for a robust data analysis are suitable procedures that facilitate the isolation of nucleic acid from microorganisms residing in complex samples, the analysis of relevant control samples, as well as high-quality genome sequence reference databases for data analysis, as exemplified in this study.

## Methods

### Vitreous Samples

A total of 21 vitreous samples from 21 individual patients were examined in this study. From April 2012 to November 2013, vitreous samples from 14 eyes with postoperative endophthalmitis following cataract surgery (n=7) and intravitreal injection (n=7) were collected using vitrectomy after informed consent had been obtained. At the Department of Ophthalmology, Glostrup Rigshospitalet (Denmark), where vitreous was collected, all patients with suspected postoperative endophthalmitis are treated with a vitrectomy independent of the presenting visual acuity. As control, vitreous was collected from 7 patients without endophthalmitis during macula hole surgery after informed consent had been obtained. Approximately 1-2 ml of vitreous body fluid was aspirated from each eye. It was at the discretion of the vitreoretinal surgeon whether to aspirate the vitreous sample before or after balanced salt solution installation. About half of each collected sample was cultured in the acute clinical setting at the Department of Microbiology, Hvidovre Hospital, Denmark, and the remaining material was stored at ‐80°C.

### Balanced salt solution samples

During vitrectomy, balanced salt solution (BSS PLUS, Alcon) is infused into the eye in order to keep the appropriate tension of the eye. BSS PLUS is a sterile physiological saltwater solution containing bicarbonate, dextrose and glutathione. Subsequently, 2.25 mg ceftazidime and 1 mg vancomycin dissolved in 0.1 ml sterile salt solution are injected into the vitreous chamber. We examined 3 paired sets of samples, i.e. 6 BSS samples in total. Aliquots were taken directly from separate BSS PLUS bottles before vitrectomy at different time points during the study period. Subsequently, BSS was collected from the vitrectomy surgical system after the BSS bottle had been connected and BSS had passed through the vitrectomy infusion line. The aliquot obtained from the vitrectomy system represents the fluid that is infused into the eye of the patient. The BSS samples were stored at ‐80°C.

### Isolation of DNA from complex samples

DNA was isolated from 200 μΐ vitreous fluid and balanced salt solution samples using two different DNA isolation procedures, i) the QIAamp DNA Mini Kit (51304, Qiagen), and ii) the QIAamp UCP Pathogen Mini Kit (50214, Qiagen). For each round of DNA isolation, one extraction control (blank) was included. For details, see Supplementary Methods.

### Metagenomic sequencing

The DNA was prepared and sequenced according to the Nextera XT DNA Library Preparation Guide, Part # 15031942 Rev. D. Sequencing was performed on an Illumina MiSeq sequencer using paired-end sequencing with v3 chemistry and 2×250 cycles. A total of 90,599,659 read pairs were obtained from the samples. The number of read pairs was in a range of 711,886 - 4,633,576 for the samples from patients with endophthalmitis, with the exception of sample I6_QIA for which only 106 read-pairs were obtained (Supplementary Table S2).

### Metagenomic sequencing data analysis

The metagenomics analysis was carried out in five steps. 1) Adapter and quality trimming as well as low complexity filtering of raw reads was performed using BBDuk of BBMap version 35.82 (http://jgi.doe.gov/data-and-tools/bbtools/). 2) Removal of human-affiliated reads from samples in a 2-step approach: i) reads that mapped against the reference genome GRCh38.p10 (GCF_000001405.36), and ii) reads that aligned to human sequences in the non-redundant nucleotide collection (nt) database from NCBI. 3) Detection of ambiguous sequences in public reference genomes and creation of curated microbial genome database that was composed of 5751 different genomes: archaea (251), bacteria (5166), fungi (225), protozoa (73), viruses (35) and the human reference GRCh38.p7 (Supplementary Table S4). 4) Classification of reads in samples using Kraken followed by Bayesian reestimation of abundance (Bracken) ^42,43^, and 5) Classification of reads using BLASTn of BLAST version 2.6.0 ^44^ For details, see Supplementary Methods.

### Cultivation and mass spectrometry (Clinical Microbiology lab)

Aliquots from the vitreous specimens were cultivated for 12 days on 5% horse blood agar, chocolate agar, brain heart infusion broth under anaerobic conditions, and on anaerobic plates (SSI Diagnostica, Denmark) under anaerobic conditions at 35°C according to the standard operating procedure at the Department of Clinical Microbiology, Hvidovre Hospital. Species identification was performed using MALDI-TOF mass spectrometry analysis (MALDI Biotyper 3.1, Bruker Daltonics Microflex LT, database MBT DB-5627) from colony material. Antimicrobial susceptibility was tested towards a range of compounds and the results were interpreted in accordance to EUCAST breakpoints (http://www.eucast.org/clinical_breakpoints/).

### Cultivation (Research lab)

To isolate bacteria and fungi from the vitreous body and balanced salt solution samples, 100 μΐ aliquots were distributed on chocolate agar (SSI Diagnostica, Denmark) and Sabouraud agar with Chloramphenicol (Fischer Scientific). The chocolate agar was incubated for 2 days at 37°C. Colonies from the chocolate agar plates were harvested and stored in Protect Multipurpose TS80 preservation tubes (Technical Service Consultants Ltd, UK) at ‐80°C. One representative colony morphotype per sample was selected for whole genome sequencing. No growth after incubation for 5 days was observed on the Sabouraud agar plates.

### Whole genome sequence analysis

Isolates were sequenced (2x150 bp paired-end) on a MiSeq system (Illumina, San Diego, CA, USA) as previously described ^45^. Reads were adapter trimmed and filtered for phiX reads using BBduk. The high-quality reads where assembled using the SPAdes assembler ^46^, and the genome sequence assemblies analysed using the Bacterial Analysis Pipeline ^47^ For details, see Supplementary Methods.

### Ethics

This study was performed in accordance with the Declaration of Helsinki. It was approved by the Danish Data Protection Agency (journal number: 2011-41-5881) and by the local ethics committee De Videnskabsetiske Komiteer - Region Hovedstaden (journal number: H-2-2011-004), and took place at public clinics in the capital region of Denmark.

### Data accessibility

The sequencing data generated and analyzed in this study are available from DDBJ/ENA/GenBank under the umbrella project PRJEB21503, including metagenomics shotgun reads (ERS1830261-ERS1830322), WGS reads (ERS1827480-ERS1827489), and WGS assemblies (ERZ468526-ERZ468535). A detailed methods description and results from the data analysis are available as supplemental material from the journal website and through Figshare (https://figshare.com/projects/Genomics- Based_Identification_of_Microorganisms_in_Human_Ocular_Body_Fluid/21038). The code for the creation of the curated microbial reference genome database is accessible from Github (https://github.com/philDTU/endoPublication), and the curated microbial reference genomes can be downloaded from ftp://ftp.cbs.dtu.dk/public//CGE/databases/CuratedGenomes.

## Acknowledgements

We thank Jacob Dyring Jensen (Technical University of Denmark) for technical assistance related to DNA sequencing. We are grateful for the support by Hanne Mordhorst (Technical University of Denmark) in the PCR-based analysis of vitreous samples, Ole Lund (Technical University of Denmark) for support with the ftp server, and Kees Veldman (Wageningen University) for providing a *S. epidermidis* reference strain. Thuy Doan (University of California San Francisco) is acknowledged for providing clarifying information regarding publication PMID: 27562436. This work was in parts supported by the European Union’s Framework Programme for Research and Innovation, Horizon2020 (643476). The funders had no role in study design, data collection and interpretation, or the decision to submit the work for publication.

## Author contributions statement

P.K., S.S.B., H.W., and S.J.P. designed the research; P.K., S.S.B., A.F.M, H.W. and S.J.P. performed the research; P.K., F.M.A., H.W., and S.J.P. contributed analytic tools; P.K., S.S.B., A.F.M., H.W., and S.J.P. analysed the data; P.K. and S.J.P. wrote the manuscript; and S.S.B, A.F.M, F.M.A., and H.W. edited the manuscript. All authors have read and approved the manuscript as submitted.

## References

1. Durand, M. L. Bacterial and Fungal Endophthalmitis. Clinical Microbiology Reviews 30, 597–613 (2017).

2. Bannerman, T. L., Rhoden, D. L., McAllister, S. K., Miller, J. M. & Wilson, L. A. The source of coagulase-negative staphylococci in the Endophthalmitis Vitrectomy Study. A comparison of eyelid and intraocular isolates using pulsed-field gel electrophoresis. Arch Ophthalmol 115, 357–361 (1997).

3. Buchta, V. et al. Outbreak of Fungal Endophthalmitis Due to Fusarium oxysporum Following Cataract Surgery. Mycopathologia 177, 115–121 (2014).

4. Foster, A. Cataract and “Vision 2020—the right to sight” initiative. British Journal of Ophthalmology 85, 635–639 (2001).

5. Solborg Bjerrum, S., Kiilgaard, J. F., Mikkelsen, K. L. & la Cour, M. Outsourced cataract surgery and postoperative endophthalmitis. Acta Ophthalmol 91, 701–708 (2013).

6. Shorstein, N. H., Winthrop, K. L. & Herrinton, L. J. Decreased postoperative endophthalmitis rate after institution of intracameral antibiotics in a Northern California eye department. Journal of Cartaract & Refractive Surgery 39, 8–14 (2013).

7. Gower, E. W. et al. Characteristics of Endophthalmitis after Cataract Surgery in the United States Medicare Population. Ophthalmology 122, 1625–1632 (2015).

8. Avery, R. L. et al. Intravitreal injection technique and monitoring: updated guidelines of an expert panel. Retina 34 Suppl 12, S1–S18 (2014).

9. McCannel, C. A. Meta-analysis of endophthalmitis after intravitreal injection of anti-vascular endothelial growth factor agents: causative organisms and possible prevention strategies. Retina 31, 654–661 (2011).

10. EndophthalmitisVitrectomyStudyGroup. Results of the Endophthalmitis Vitrectomy Study. A randomized trial of immediate vitrectomy and of intravenous antibiotics for the treatment of postoperative bacterial endophthalmitis. Endophthalmitis Vitrectomy Study Group. Arch Ophthalmol 113, 1479–1496 (1995).

11. Yao, K. et al. The incidence of postoperative endophthalmitis after cataract surgery in China: a multicenter investigation of 2006-2011. British Journal of Ophthalmology 97, 1312–1317 (2013).

12. Seal, D. et al. Laboratory diagnosis of endophthalmitis: Comparison of microbiology and molecular methods in the European Society of Cataract & Refractive Surgeons multicenter study and susceptibility testing. Journal of Cataract & Refractive Surgery 34, 1439–1450 (2008).

13. Barry, P., Cordovés, L. & Gardner, S. ESCRS guidelines for prevention and treatment of endophthalmitis following cataract surgery: data, dilemmas and conclusions. (ESCRS, 2013).

14. Wilson, M. R. et al. Actionable Diagnosis of Neuroleptospirosis by Next-Generation Sequencing. New England Journal of Medicine 370, 2408–2417 (2014).

15. Oh, J. et al. Temporal Stability of the Human Skin Microbiome. Cell 165, 854–866 (2016).

16. Zankari, E. et al. Identification of acquired antimicrobial resistance genes. Journal of Antimicrobial Chemotherapy 67, 2640–2644 (2012).

17. Joensen, K. G. et al. Real-Time Whole-Genome Sequencing for Routine Typing, Surveillance, and Outbreak Detection of Verotoxigenic Escherichia coli. Journal of Clinical Microbiology 52, 1501–1510 (2014).

18. Breitwieser, F. P. & Salzberg, S. L. Pavian: Interactive analysis of metagenomics data for microbiomics and pathogen identification. (bioRxiv, 2016). doi:10.1101/084715

19. Doan, T. et al. Illuminating uveitis: metagenomic deep sequencing identifies common and rare pathogens. Genome Medicine 1–9 (2016). doi:10.1186/s13073-016-0344-6

20. Mattos, F. B., Saraiva, F. P., Angotti-Neto, H. & Passos, A. F. Outbreak of Ochrobactrum anthropi endophthalmitis following cataract surgery. Journal of Hospital Infection 83, 337–340 (2013).

21. Venkateswaran, N., Wozniak, R. A. F. & Hindman, H. B. Ochrobactrum anthropi Keratitis with Focal Descemet's Membrane Detachment and Intracorneal Hypopyon. Case Reports in Ophthalmological Medicine 2016, 1–4 (2016).

22. Relman, D. A. Actionable Sequence Data on Infectious Diseases in the Clinical Workplace. Clinical Chemistry 61, 38–40 (2014).

23. Pallen, M. J. Diagnostic metagenomics: potential applications to bacterial, viral and parasitic infections. Parasitology 141, 1856–1862 (2014).

24. Didelot, X., Bowden, R., Wilson, D. J., Peto, T. E. A. & Crook, D. W. Transforming clinical microbiology with bacterial genome sequencing. Nature Reviews Genetics 13, 601–612 (2012).

25. Mollerup, S. et al. Propionibacterium acnes: Disease-Causing Agent or Common Contaminant? Detection in Diverse Patient Samples by Next-Generation Sequencing. Journal of Clinical Microbiology 54, 980–987 (2016).

26. Salter, S. J. et al. Reagent and laboratory contamination can critically impact sequence-based microbiome analyses. BMC Biology 1–12 (2014). doi:10.1186/s12915-014-0087-z

27. Glassing, A., Dowd, S. E., Galandiuk, S., Davis, B. & Chiodini, R. J. Inherent bacterial DNA contaminationof extraction and sequencing reagents may affect interpretation of microbiota in low bacterial biomass samples. Gut Pathogens 1–12 (2016). doi:10.1186/s13099-016-0103-7

28. Tanner, M. A., Goebel, B. M., Dojka, M. A. & Pace, N. R. Specific ribosomal DNA sequences from diverse environmental settings correlate with experimental contaminants. Applied and Environmental Microbiology 64, 3110–3113 (1998).

29. Naccache, S. N. et al. The Perils of Pathogen Discovery: Origin of a Novel Parvovirus-Like Hybrid Genome Traced to Nucleic Acid Extraction Spin Columns. Journal of Virology 87, 11966–11977 (2013).

30. Smuts, H., Kew, M., Khan, A. & Korsman, S. Novel hybrid parvovirus-like virus, NIHCQV/PHV, contaminants in silica column-based nucleic acid extraction kits. Journal of Virology 110, 10264–10269 (2014).

31. Lee, A. Y., Akileswaran, L., Tibbetts, M. D., Garg, S. J. & Van Gelder, R. N. Identification of Torque Teno Virus in Culture-Negative Endophthalmitis by Representational Deep DNA Sequencing. Ophthalmology 122, 524–530 (2015).

32. Merchant, S., Wood, D. E. & Salzberg, S. L. Unexpected cross-species contamination in genome sequencing projects. PeerJ 2, e675 (2014).

33. Kryukov, K. & Imanishi, T. Human Contamination in Public Genome Assemblies. PLoS ONE 11, e0162424 (2016).

34. Longo, M. S., O'Neill, M. J. & O'Neill, R. J. Abundant Human DNA Contamination Identified in Non-Primate Genome Databases. PLoS ONE 6, e16410 (2011).

35. Harper, D. R. A comparative study of the microbiological contamination of postmortem blood and vitreous humour samples taken for ethanol determination. Forensic Sci Int 43, 37–44 (1989).

36. Egger, S. F. et al. Bacterial growth in human vitreous humor. Exp Eye Res 65, 791–795 (1997).

37. Hasman, H. et al. Rapid Whole-Genome Sequencing for Detection and Characterization of Microorganisms Directly from Clinical Samples. Journal of Clinical Microbiology 52, 139–146 (2014).

38. Joensen, K. G. et al. Evaluating next-generation sequencing for direct clinical diagnostics in diarrhoeal disease. European Journal of Clinical Microbiology & Infectious Diseases 36, 1325–1338 (2017).

39. Greninger, A. L. et al. Rapid metagenomic identification of viral pathogens in clinical samples by real-time nanopore sequencing analysis. Genome Medicine 1–13 (2015). doi:10.1186/s13073-015-0220-9

40. Cao, M. D. et al. Streaming algorithms for identification of pathogens and antibiotic resistance potential from real-time MinIONTM sequencing. GigaScience 5, 32 (2016).

41. Quick, J. et al. Multiplex PCR method for MinION and Illumina sequencing of Zika and other virus genomes directly from clinical samples. Nat Protoc 12, 1261–1276 (2017).

42. Wood, D. E. & Salzberg, S. L. Kraken: ultrafast metagenomic sequence classification using exact alignments. Genome Biology 15, 1–12 (2014).

43. Lu, J., Breitwieser, F. P., Thielen, P. & Salzberg, S. L. Bracken: estimating species abundance in metagenomics data. PeerJ Computer Science 3, e104 (2017).

44. Altschul, S. F., Gish, W., Miller, W., Myers, E. W. & Lipman, D. J. Basic local alignment search tool. J Mol Biol 215, 403–410 (1990).

45. Bartels, M. D. et al. Comparing Whole-Genome Sequencing with Sanger Sequencing for spa Typing of Methicillin-Resistant Staphylococcus aureus. Journal of Clinical Microbiology 52, 4305–4308 (2014).

46. Bankevich, A. et al. SPAdes: A New Genome Assembly Algorithm and Its Applications to Single-Cell Sequencing. Journal of Computational Biology 19, 455–477 (2012).

47. Thomsen, M. C. F. et al. A Bacterial Analysis Platform: An Integrated System for Analysing Bacterial Whole Genome Sequencing Data for Clinical Diagnostics and Surveillance. PLoS ONE 11, e0157718 (2016).

